# Prioritized adjustments in posture stabilization and adaptive reaching during neuromuscular fatigue of lower-limb muscles

**DOI:** 10.1101/2024.04.02.587637

**Authors:** Mauro Nardon, Oindrila Sinha, John Kpankpa, Eliza Albenze, Cédrick Bonnet, Matteo Bertucco, Tarkeshwar Singh

## Abstract

Neuromuscular fatigue (NMF) induces temporary reductions in muscle force production capacity, affecting various aspects of motor function. While studies have extensively explored NMF’s impact on muscle activation patterns and postural stability, its influence on motor adaptation processes remains less understood. This paper investigates the effects of localized NMF on motor adaptation during upright stance, focusing on reaching tasks. Utilizing a force field perturbation paradigm, participants performed reaching movements while standing upright before and after inducing NMF in the ankle dorsiflexor muscles. Results revealed that despite maintained postural stability, participants in the NMF group exhibited larger movement errors during reaching tasks, suggesting impaired motor adaptation. This was evident in both initial and terminal phases of adaptation, indicating a disruption in learning processes rather than a decreased adaptation rate. Analysis of electromyography activation patterns highlighted distinct strategies between groups, with the NMF group showing altered activation of both fatigued and non-fatigued muscles. Additionally, differences in co-activation patterns suggested compensatory mechanisms to prioritize postural stability despite NMF-induced disruptions. These findings underscore the complex interplay between NMF, motor adaptation, and postural control, suggesting a potential role for central nervous system mechanisms in mediating adaptation processes. Understanding these mechanisms has implications for sports performance, rehabilitation, and motor skill acquisition, where NMF may impact the learning and retention of motor tasks. Further research is warranted to elucidate the transient or long-term effects of NMF on motor adaptation and its implications for motor rehabilitation interventions.

**New & Noteworthy:** We assessed motor adaptation during force field reaching following exercise-induced neuromuscular fatigue (NMF) on postural muscles. NMF impaired adaptation in performance. Similarly, diverging activation strategies were observed in the muscles. No effects were seen on measures of postural control. These results suggest the remodulation of motor commands to the muscles in presence of NMF, which may be relevant in settings where participants could be exposed to NMF while learning, such as sports and rehabilitation.

## Introduction

Exercise-induced neuromuscular fatigue (NMF) is characterized by a temporary reduction in the ability of a muscle to produce force and/or power, (1–3). This phenomenon affects various dimensions of human motor function, including the planning of motor activities (4–7), coordination (8–11), balance (12), sensorimotor integration (13, 14), limb proprioception (15, 16) and muscle activation patterns (17). Furthermore, NMF adversely affects gait characteristics (18–21) and posture stabilization (22–24). Specifically, NMF leads to postural instability by enhancing the sway velocity of the center of pressure (22) and diminishing the efficacy of balance recovery following disturbances (25). Contrarily, some investigations have reported no direct impact of NMF on postural stability metrics, despite noting variations in muscle activation patterns (17, 26). These alterations in muscle activation post-NMF are interpreted by subsequent analyses as efficient adaptations by the central nervous system (CNS) to counteract NMF and preserve postural steadiness (4, 17, 23, 24, 26).

Exercise-induced unilateral NMF induces central adaptations that manifest as adaptive (and sometimes maladaptive) adjustments in the motor output of the contralateral, non-exercised limb (27, 28). This phenomenon, known as non-local muscle fatigue, involves performance alterations in contralateral or distant, non-exercised muscle groups. It is primarily mediated by central mechanisms and is believed to involve neural structures responsible for coordinating bilateral and unilateral movements between the upper and lower extremities (29). Notably, the cerebellum, which plays a crucial role in the integration of proprioceptive inputs for the real-time and offline control of movements, may also support fatigue induced central adaptations in the descending drive to exercised and unexercised muscles. Indeed, recent evidence indicates that a reduction in cerebellar excitability is associated with an altered perception of fatigue and changes during visuomotor adaptation (30). Considering the cerebellum’s role in coordinating bilateral movements (31, 32), and integrating motor functions between the upper and lower body (33) it is conceivable that NMF significantly impacts motor functions through cerebellar mediation, such as during motor adaptation tasks.

The motor adaptation paradigm, which investigates the sensorimotor system’s ability to recalibrate in response to experimental perturbations, has been instrumental in establishing the cerebellum as a critical component of the neural circuitry facilitating adaptation (34–38). Within this paradigm, the force field adaptation approach involves participants executing reaching movements under conditions of artificially induced perturbations, such as velocity-dependent viscous or curl force fields generated by robots, thereby learning to recalibrate their motor output in response to altered proprioceptive feedback on a trial-by-trial basis (39).

A study by Takahashi and colleagues (40) revealed that NMF can impair the development and retention of a recalibrated mapping between altered proprioceptive feedback and motor output necessary to perform a force field reaching task. This finding implies that NMF may specifically influence adaptation in unilateral upper-extremity tasks, but it also raises questions about whether such interference is localized to the limb engaged in the exercise or if adaptive (or maladaptive) modifications extend to non-exercised limbs, thereby affecting overall performance and coordination. Addressing this question would enhance our comprehension of the mechanisms by which the nervous system concurrently maintains global postural stability and facilitates movement through localized activation of upper limb musculature.

A simple approach to explore this question would involve conducting a force field reaching task during upright stance. Ahmed and Wolpert (41) utilized this paradigm and compared the transferability of learned novel force field dynamics between seated and upright standing conditions. Their findings demonstrated that participants not only learned the novel dynamics but also successfully generalized these learnings across different postural stances (from seated to standing and vice versa). Additionally, from the onset of trials, individuals exhibited the capability for making pertinent anticipatory postural adjustments while standing. Collectively, these results imply that the networks responsible for posture stabilization and movement coordination can concurrently manage different motor functions to adeptly navigate actions within new dynamic environments.

Based on these disparate sets of studies, we hypothesized that the nervous system maintains postural stability and movement accuracy during NMF by modulating the central descending drive to muscles. During fatigue, earlier anticipatory postural adjustments (APAs) indicate a functional adaptation by the motor system to maintain postural stability (7). Studies have shown that fatigue degrades proprioception in the fatigued muscles, leading the nervous system to likely reweight sensory information from these muscles (42) and increase the descending drive to the non-fatigued muscles (43). These adjustments reflect central compensation and task prioritization to stabilize posture when fatigue alters the peripheral sensorimotor state.

Therefore, when leg muscles are fatigued, we expect the descending drive to other lower extremity muscles to be modulated to stabilize posture (2, 44). Exercise-induced fatigue also alters neural activity in areas like the cerebellum and frontal cortical regions, which are involved in both postural balance and goal-directed reaching movements (45–48). These areas would likely modulate the descending drive to lower extremity muscles to stabilize posture and maintain motor performance. Based on this hypothesis, we predict alterations in the activation patterns of both exercised and unexercised muscles to maintain reaching accuracy and stabilize center-of-pressure dynamics during a force field adaptation task. We tested this prediction using an experimental task where participants performed reaching movements under a force field while standing upright.

## Materials and Methods

### Participants

Twenty-eight right-handed, healthy young participants (14 males; 14 females) were randomly assigned to either a *Fatigue* or *No-Fatigue* group. Participants were excluded if they had a history of neurological or musculoskeletal disorders, cardiovascular disease that would make it difficult to perform the exercises, and conditions of the vestibular system that would affect balance. Participants were also excluded if they took medications which made them drowsy and would affect upright posture control. The study protocol conformed to the most recent principles of the Declaration of Helsinki and was approved by the Institutional Review Board of The Pennsylvania State University. Participants provided written informed consent before taking part in the study. Participants’ demographics are reported in ***Table 1***.

### Apparatus

Experiments were carried out employing a unimanual *KINARM* End-Point robot (Kinarm Inc, Kingston, Canada), in an adjustable height configuration for conducting reaching studies during upright stance. The system was integrated with a force plate (FP4060-07-TM, Bertec, Columbus, OH, USA) positioned in front of the center of the robot screen. Participants’ safety was ensured by using a full-body harness, connected by carabiners to an overhead metal rack (max operating weight and height range: 163 kg and 97-213 cm, respectively; **Figure 1** A).

**Figure 1.**
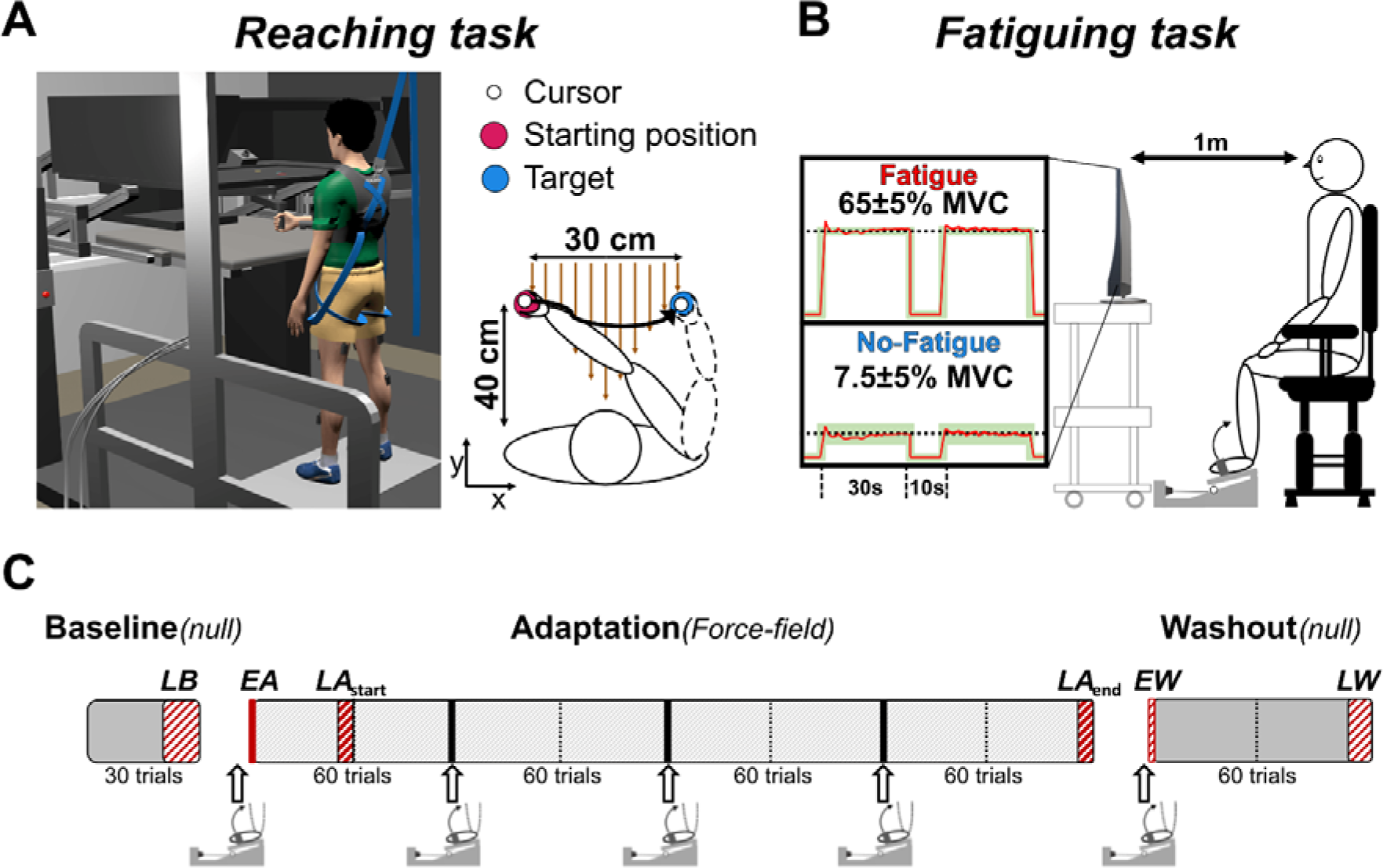
Schematic representation of the experimental setup. A: experimental setup (Side view) of the reaching task on the KINARM robot screen (Top view). Brown, vertical arrows represent the direction of force field perturbation, while the black solid line represents the cursor trajectory during a reaching movement. Laboratory axes orientation is shown at the bottom-left corner. B: illustration for the isometric ankle dorsiflexion exercise setup. Knee and ankle joint angles were kept ∼90° and 110° throughout the exercise. Real-time visual feedback of the force was provided by a monitor, which also displayed a green-shaded area corresponding to the required force target differing between groups (Fatigue= 65 ± 5%, No-Fatigue= 7.5 ± 5% MVC). C: schematic outline of the experimental phases: baseline phase consisted of 30 unperturbed (null) reaching trials, adaptation phase consisted of 240 perturbed (force field) trials divided into 4 blocks of 60 trials and washout phase consisted of 60 null trials. Bottom isometric setup-like symbols indicate the fatiguing exercise bouts (see Isometric Task in the text). Thin vertical dotted lines separating adaptation and washout blocks represent short breaks (<1min) in between two blocks of 30 trials during the adaptation and washout phases (see Reaching Task in the text). Experimental phases considered for the analyses are highlighted in red. LB: late baseline, EA: early adaptation, LA_start_: late adaptation – start, LA_end_: late adaptation – final block, EW: early washout, LW: late washout.

Electromyographic (EMG) activity of eleven muscles – three in the upper right limb (*anterior deltoid* (AD), *posterior deltoid* (PD), and *triceps brachii* (TR) and four each in both the lower limbs, bilaterally (*rectus femoris* (RF), *biceps femoris* – long head (BF), *tibialis anterior* (TA), *gastrocnemius medialis* (GM) – was recorded using wireless EMG sensors (Trigno, Delsys Inc., Natick, MA, USA). Sensor positioning was conducted following recommendations (49), signal baseline noise and signal-to-noise ratio were inspected using the dedicated software (*EMGWorks*, Delsys Inc.) to ensure high EMG signal quality. EMG signals were digitized and synchronized with robot kinetics and kinematics, and with force-plate data through an A/D converter (National Instruments) and automatically streamed to the computer running the robot data-acquisition software (Dexterit-E™, Kinarm Inc, Kingston, ON, Canada). Data from the robotic manipulandum, force plate and EMG were synchronized and sampled at 1000 Hz.

A customized frame equipped with a uniaxial strain-gauge force sensor (Model S-AL-A, Deltatech, Sogliano al Rubicone, Italy) was used for the force measurement during the isometric fatiguing task (see *Procedures*) to measure isometric force in ankle dorsiflexion. Force data were collected using a custom LabVIEW VI (National Instruments) software at 50 Hz sampling rate. Signals were exported and analyzed offline using MATLAB (R2023a, version 9.14.0, MathWorks, Natick, MA, USA).

### Procedures

Participants were given an overview of the experiment and received specific instructions: 1) avoid leaning the forehead on the Kinarm while standing; and 2) avoid pulling the robot handle downward. They first performed 10 (unperturbed) trials to familiarize with the reaching task. Few additional isometric trials (*normTrials*) were performed to be used for EMG normalization.

### Reaching task

Participants stood barefoot on the force plate, in front of the robot screen. Feet positions were checked and marked on the force platform to ensure consistency across experimental blocks. KINARM screen and handle height was adjusted to each participant’s height to ensure comfortable range of movement. The task consisted of planar reaching movements in the medio-lateral direction (range: 30 cm). Participants’ vision of the hand was occluded by the screen during the trials and they were required to move the cursor (radius: 1 cm) – corresponding to the hand position – to the location of the targets on the screen. Targets were within reach of the participant and did not require trunk movement. The starting position consisted of a circle (radius: 1.5 cm; positioned 5 cm left from the mid-sagittal plane, ∼40 cm away from participant’s chest). Once reached, the data acquisition started, and participants were instructed to maintain the cursor in this location until a second target (radius: 1.5 cm) appeared 30 cm to the right (**Figure 1** A) and reach for it as quickly and accurately as possible. Targets color changed when reached by the participant. The experiment was programmed to display the target with a fixed (1,500 ms) plus a random delay (1-500 ms) after the starting position had been reached to avoid predictability of the stimulus and its anticipation. If participants moved the cursor before the target appeared, the trial was aborted and repeated. Participants were instructed to return to *start* at a comfortable pace. Immediately after familiarization, participants performed a block (*baseline*; 30 trials) of unperturbed (*null*) reaching movements. During the subsequent experimental phase (*adaptation;* 240 trials), the robot simulated a viscous curl field (*force field*) by generating a force (*F*) perpendicular (clockwise) to the direction and proportional to the magnitude of the instantaneous velocity (*V*) of the robot handle:

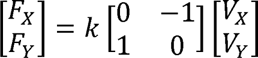

Where the field gain *k* was set to 0.2 N*s/cm. The last experimental phase (*washout*; 60 trials) consisted of unperturbed (*null*) trials. In order to avoid fatigue in the upper limb, participants were allowed short breaks (<1min) to relax their arm and posture halfway (every 30 trials) in *adaptation* and *washout* blocks. This was based on pilot work conducted in the laboratory prior to the study. Handle position, velocity and the force applied at the handle, ground reaction forces, center of pressure (CoP) and surface EMG data were recorded during each trial.

### Isometric fatiguing task

Participants seated on a height-adjustable chair with their feet fastened to a customized metal frame through two inextensible straps, placed proximal to the metatarsophalangeal joints (**Figure 1** B). Knee and ankle joint angles were maintained at approximately 90° and 110°, respectively and the position did not change throughout the entire experimental session. To avoid the involvement of other postural and upper limb muscles, participants were instructed to place and keep their hands prone on their thighs.

Participants performed three ∼5 s maximal voluntary contractions (MVCs) in isometric dorsiflexion of the ankles, using mainly both their tibialis anterior muscles. MVCs were inspected and the highest value was used to set the relative exercise intensity. To ensure maximal effort and avoid errors in the MVC determination, if one of the MVCs value differed more than 10% from the others, the value was discarded and an additional MVC trial was performed. Participants rested for >1 min between consecutive MVCs.

Sustained, cyclical isometric exercise protocol (40 seconds; 75% duty cycle) was set relative to each participant’s MVC value and differed between groups (Fatigue: 65 ± 5%; No-Fatigue: 7.5 ± 5% MVC) (50). This ensured that both groups went through identical experimental protocols, but only one group was fatigued. This resulted with induced localized neuromuscular muscle fatigue on the main ankle dorsiflexor muscles – TAs for the Fatigue group. A computer monitor (22”; viewable diagonal screen size 54.8 cm), placed ∼1 m away at participant’s eye level, provided real-time visual feedback on the force production during the exercise while an audible timer set the pace of the exercise-rest phases of the protocol. Participants were asked to keep the force – shown as a black dot – within the target force boundaries, displayed on the monitor screen as a green-shaded area throughout the entire exercise phase (30s) of the cycle and successively relax for the remaining 10s of the cycle.

Participants in the Fatigue group were considered fatigued and stopped when their force did not reach and sustain for >1s the target force for two consecutive cycles despite continuous encouragement. Few seconds later, participants were asked to produce an additional MVC and then asked to move back to the reaching task and immediately perform the subsequent movement block (≤30 s).

Mean exercise time for each bout (each exercise in between blocks of reaching task) of isometric tasks of the Fatigue group was calculated and rounded up to the nearest cycle (40s). These exercise times were later used to stop participants in the No-Fatigue group (exercising around 7.5% MVC). This was done to ensure a similar time duration in both groups and required the entire Fatigue group to be tested before No-Fatigue. Participants in both groups were stopped by the experimenter and were blind to exercise intensity (%MVC – i.e. y-axis values were hidden from the feedback graph), termination criteria and exercise bouts duration. The isometric exercise setup was positioned ∼1.5 meters away from the robot to ensure a rapid transition between the tasks.

Since NMF is a transient phenomenon (10, 51, 52), fatiguing exercise was repeated multiple times throughout the experiment to limit recovery processes. To assess the effects of NMF on motor adaptation, we computed performance variables, postural stability measures and EMG activity during anticipatory (AP) and corrective windows – here we define EMG activity occurring after movement initiation as early- and voluntary Reactive Responses (eRR and vRR, respectively).

### Data analysis

Computed variables were compared at six phases of the experiment: *late baseline* (LB; last 10 trials in baseline phase); *early adaptation* (EA; 1^st^ trial in *adaptation* phase); *late adaptation* – *start* (LA_start_; 27^th^–30^th^ trial); *late adaptation* – *final* (LA_end_; last 5 trials in *adaptation* phase); *early washout* (EW; 1^st^ trial in *washout* phase); *late washout* (LW; last 5 trials in *washout* phase). Separate planned comparisons were performed across phases: 1) LB, EA, LA_start_, LA_end_; and 2) LB, EW, LW, to assess adaptation rate and *washout* phase effects, respectively, in accordance with previous similar studies (41, 53, 54). A schematic outline of the experiment and the relative phases is shown in **Figure 1** (C).

Handle kinematics, hand applied force, and force plate signals were filtered with a 10 Hz low-pass, 6^th^ order, zero-phase digital Butterworth filter. EMG signals were detrended, band-pass filtered with a 20–450 Hz, 4^th^ order, zero-phase digital Butterworth filter and rectified, then a 57–63 Hz, band-stop, 4^th^ order filter was applied to remove line noise. Hand movement onset (*t0*) was calculated as the point in time when hand force profile in the mediolateral direction crossed 5% of its peak value (similar to (55) - **Figure 2**). EMG, hand kinematics and force plate signals were then aligned from 500 ms before-to 1,000 ms after *t0* and stored for further analyses. EMG signals from *normTrials* were similarly pre-processed and stored separately. Reaction time (the time elapsed between target presentation and *t0*) and movement time (the time between *t0* and the instance when the cursor reached *Target*) were computed to control for variation in movement parameters.

**Figure 2.**
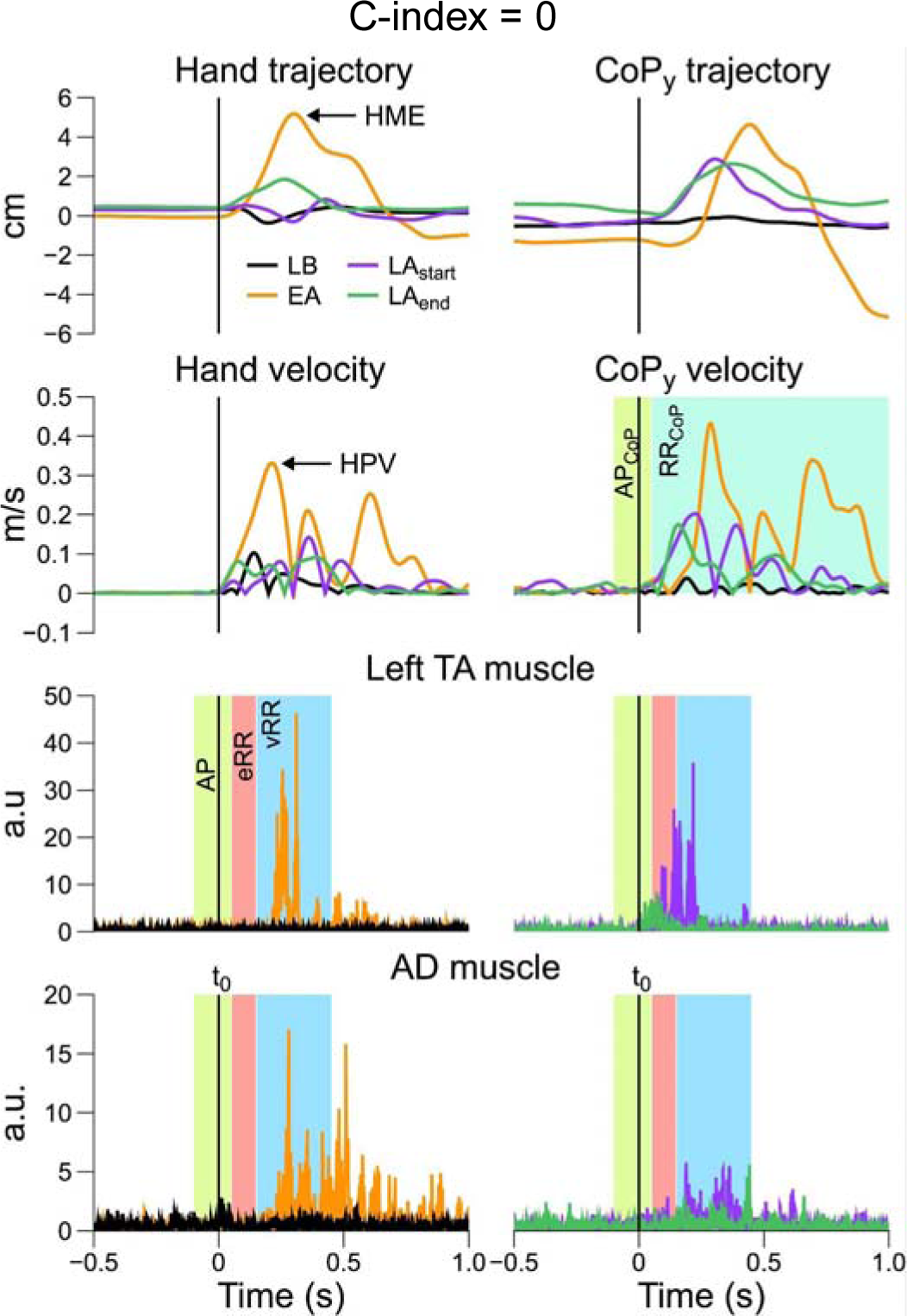
Aligned signals for a representative participant across different experimental phases (different colors - see legend). Vertical black line (t0) corresponds to the hand movement onset. Shaded areas highlight CoP-based and EMG-based calculation windows (see: Data analysis in text). AD: anterior deltoid, AP: anticipatory phase, EA: early adaptation, eRR: early Reactive Response, EW: early washout, HME: hand movement error, HPV: hand peak velocity, LA_end_: late adaptation – final block, LA_start_: late adaptation – start, LB: late baseline, LW: late washout, RR: reactive response, TA: tibialis anterior, vRR: voluntary Reactive Response.

### Measures of overall performance

Hand movement error (HME) was calculated as the peak signed perpendicular deviation of the hand trajectory from a straight line connecting the two targets (41). Positive values represent an error in the backward direction (towards the participant’s body). As a measure of hand trajectory corrections, absolute peak hand velocity in the anteroposterior direction (HPV) was computed during the movement phase. Signal profiles for a representative participant are shown in **Figure 2** (Left panels - 1^st^ and 2^nd^ line). We performed an additional analysis on the adaptation profiles and adaptation rates of the two groups by considering HME and HPV values during the first block of trials during the adaptation phase (n=54 trials). We chose to fit only the first block of trials, since HME and HPV values reduced rapidly, with values that tend to stabilize after ∼40-50 trials. Following each bout of fatiguing exercise, participants showed a slight increase (i.e. sort of forgetting effect) in HME and HPV values for the first trials during re-exposure to perturbation. Individual data were smoothed using a 9-point moving average filter and then averaged within each group, similarly to Malone et al., (56). A single-term exponential function was then fitted to each resulting dataset using the following formula in MATLAB:

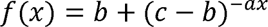

Fitting coefficients were then compared between groups and considered different when their 95% confidence intervals did not overlap.

### Measures of postural stability

Center of pressure (CoP) position was computed from force plate data according to Winter (57). CoP position was then demeaned using the mean value of CoP during the first 5 samples of the trial. Maximal backward and forward displacement of the CoP during movement (CoP_back_ and CoP_forward_, respectively) were calculated as the lowest or highest CoP value in the anteroposterior direction (CoP_y_ trajectory, where negative values represent backward displacement from initial position). CoP anticipatory phase (AP_CoP_) was quantified as the mean CoP velocity in the anteroposterior direction (CoP_y_ velocity) over a 150 ms time window starting 100 ms prior- and ending 50 ms after movement onset (*t0*). CoP reactive response (RR_CoP_) – considered an index of postural corrections (41) – was calculated as the maximum absolute value of CoP_y_ velocity over the rest of the movement (from 50ms after *t0* until the end of the trial). Calculation windows for AP_CoP_ and RR_CoP_ are shown in the upper right panels of **Figure 2**. Time-windows were based on earlier work (58) and consistent with previous motor adaptation studies (41).

### Measures of muscle activation

EMG median frequency of both *tibialis anterior* muscles during the fatiguing task was computed from pre-processed signals and then averaged between the two muscles in order to confirm fatigue-induced changes in muscle spectral power frequency following the fatiguing protocol.

Linear envelopes of pre-processed EMG signals were computed by applying a 100 Hz low-pass, 4^th^ order, zero-phase digital Butterworth filter. The resulting aligned linear envelopes were then averaged participant-wise across the first/last three consecutive trials for each experimental phase (59), respectively (e.g., for *late baseline* (LB): average of last 3 trials; for *early adaptation* (EA): average of first 3 trials). Concerning *normTrials –* 10 trials where participants had to maintain the hand cursor over a circle-shaped target (1.5 cm radius) and a background constant load of –10 N was applied in the backward direction to activate the muscles – the relative EMG envelopes were averaged across all the trials, yielding a single averaged signal for each muscle.

Phase-averaged EMG signals were then integrated at the level of each muscle using a trapezoidal numerical integration (function “*trapz*” in MATLAB software) for three pre-defined time windows, relative to *t0*: *Anticipatory Phase* (AP) from –150 to +50 ms, *early Reactive Response* (eRR) over a 100 ms window from +50 to +150 ms, and *voluntary Reactive Responses* (vRR) over a 300 ms window from +150 to +450 ms. Time-windows for EMGs integration were chosen based on the nature of the reaching task and on the continuous nature of the force field perturbation. The lower panels of **Figure 2** show a representative muscle in the lower limb (Left TA) and one in the upper limb (AD), respectively. Reactive responses windows differentiate on the physiological nature of the activity: the first window (eRR) mainly represents spinal and supraspinal stretch reflexes responses (60), while vRR conveys information about voluntary, visually-driven EMG activity ensuring online trajectory correction, which are characterized by longer latencies (∼120 to 200 ms in the lower limbs), as shown in previous work (61).

Computed integrals were further normalized for each participant by dividing it by the integral of the averaged EMG activity during *normTrials* (∫norm*EMG*; integrated over a 100 ms time-window, starting 100 ms after the force was applied).

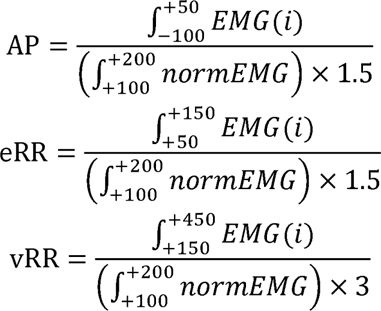

We further normalized EMG activity participant-wise by dividing integrated EMG values of each time-window (i.e. AP, eRRs and vRRs) for each experimental phases (i.e. EA, *LA_start_*, *LA_end_*, EW, LW) by the relative integrated EMG during late baseline (LB) – similarly to (54) – to control for baseline differences and explore changes in muscle activation across phases during APs, eRRs and vRRs.

Finally, we computed indexes of co-activation (C-index) within agonist–antagonist muscle pairs acting at joint level (Left TA/GM, Right TA/GM, Left RF/BF, Right RF/BF and AD/PD), using EMG integrals of ventral and dorsal muscles (before normalization by *late baseline*) for each time-epoch (62–66). Specifically:

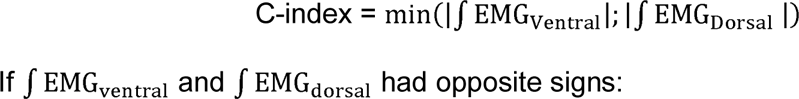

### Statistical analysis

Jamovi software (Version 2.3.28; The Jamovi project (2023); https://www.jamovi.org) was used for statistical analyses. All data in the text, Tables, and Figures – unless otherwise stated – are presented as mean ± 1 standard deviation (SD). For all comparisons, statistical significance was set at an α of 0.05. Data distribution (skewness and kurtosis) was assessed, the normality of the data was assessed by the Shapiro–Wilk test and confirmed by the inspection of density and Q–Q plots. When normality assumption was violated, non-parametric statistical tests were performed (Friedman’s test with pairwise Durbin-Conover comparisons). Successively, independent samples t-test (Welch-test) was used for pairwise comparisons between groups.

To test the effects of the fatiguing isometric exercise protocol, group differences in MVC changes before and at the end of each bout were assessed using a two-way repeated measures ANOVA (*Group* X *Time* – with *Time* as a repeated measure factor). Paired samples t-tests were performed to compare EMG median frequency of *tibialis anterior* muscles during the 1^st^ and the last cycle of the 1^st^ bout of isometric exercise in the fatiguing protocol, for each group separately.

To test the effects of fatigue on the adaptation process, a two-way repeated measures ANOVA *Group* (Fatigue and No-Fatigue) X *Phase* (LB, EA, LA_start_ and LA_end_ – with *Phase* as a repeated measure factor) was used to assess group differences in performance and postural variables across four experimental phases (LB, EA, LA_start_, LA_end_). Assumptions of sphericity were explored using Mauchly’s test and controlled for using the Greenhouse–Geisser adjustment in instances where Mauchly’s test was significant (α <0.05). The corrected degrees of freedom are reported in results. In the event of a significant interaction or main effect, post hoc comparisons were performed using Holm-Bonferroni correction. Independent-sample t-tests (Welch-test) were used to assess between-group differences if Levene’s test revealed unequal variance.

Similarly, to test the effects of fatigue on the washout process, a two-way repeated measures ANOVA (*Group* X *Phase* – *Phase* as a repeated measure factor) was used to assess group differences in performance and postural variables across three experimental phases (LB, EW, LW). Integrated EMG time-windows were compared between groups using a two-way repeated measures ANOVA (*Group* X *Phase – Phase* as a repeated measure factor) where *Phase* levels were N-1 due to the normalization to LB values (*Phase* as a repeated measure factor). To test for differences within each group from LB values, a one-way repeated measure ANOVA (*Phase* as a repeated measure factor) was performed for each group (Fatigue/No-Fatigue), separately.

Similar to Mathew et al. (67), for CoP-related variables – since our initial hypothesis assumed the equality (i.e. no differences) of these variables between groups – a Bayesian approach (Bayesian rm ANOVA) on JASP software (Version 0.18.3; JASP Team (2024)) was used to confirm the statistical equivalence in cases where the *null* hypothesis for the factor *Group* was not rejected by traditional ANOVA. In such cases we calculated Bayes’ factors, which quantify the degree to which the data favors either one or the other model (i.e. means are equals vs means are different –(68–70)). According to this definition, *BF_01_* is the evidence supporting H0 (i.e. means are equals), which was conventionally interpreted according to Kass and Raftery (71): *BF_01_* between 1 and 3: barely worth mentioning; *BF_01_* between 3 and 20: positive; *BF_01_* > between 20 and 150: strong; *BF_01_* > 150: very strong evidence.

## Results

Participants adapted throughout the trials to the introduction of the novel perturbation, both in terms of hand trajectory and in terms of postural measures and muscle activation. As expected, participants in both groups showed appreciable improvement in the reaching task performance, with adaptation strategies persisting even after the removal of the force field perturbation, as shown by the aftereffects – opposite in direction – on performance in the washout phase (**Figure 3**). NMF caused larger hand movement errors (HME; **Figure 3**) during the adaptation phase for the *Fatigue* group. There were also between-group differences for hand trajectory corrections estimated from hand peak velocity (HPV; **Figure 3**). The alterations in hand trajectory do not support our prediction that reaching movements would be similar between the *Fatigue* and *No-Fatigue* groups.

**Figure 3.**
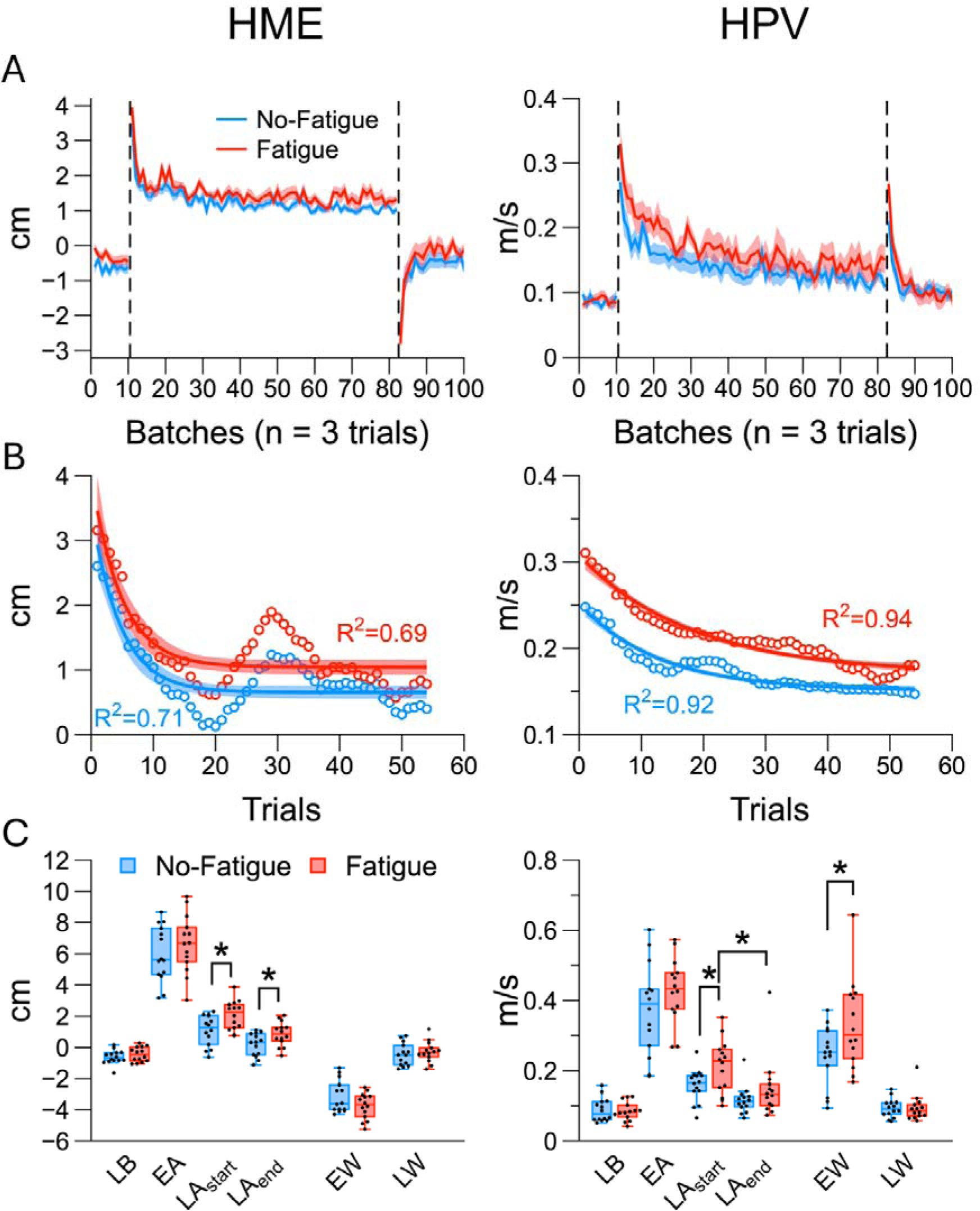
Overall performance variables during the reaching task divided by group. A) Batch-averaged curves for hand movement error (HME; left panel) and hand peak velocity (HPV; right panel) throughout the experimental phases. Positive HME values represent a backward error (towards the participant). HPV was computed in absolute values. Vertical dotted lines represent the transition across experimental phases (baseline, adaptation, washout). Data are averaged in batches of n=3 trials and reported as mean (thick lines) ± 1 Standard Error (SE; shaded areas). B) Fitting curves (thick lines) and confidence intervals (C.I.) of the fittings (shaded areas) for HME (left) and HPV (right) group-averaged values (circles) during the first block of adaptation trials. C) Box plots for HME (left) and HPV (right) values across the considered experimental phases. Boxes depict data from first to third quartile, whiskers represent minimum and maximum data points. Median is shown by the horizontal line. * indicates significant effects (p < 0.05) between groups. EA: early adaptation, EW: early washout, LA_end_: late adaptation – final block, LA_start_: late adaptation – start, LB: late baseline, LW: late washout.

However, our second prediction was confirmed: measures of postural stability were similar between the two groups (**Figure 6**). This was likely due to fatigue-induced changes in postural muscle activation in multiple muscles (**Figures 4 & 5**). Between-group differences in muscle activation were not limited to the fatigued muscles, but extended to others, non-fatigued muscles and even to upper-limb muscles. These results suggest that during fatigue, the nervous system prioritizes the stabilization of posture through changes in the descending drive to both lower and upper extremity muscles.

**Figure 4.**
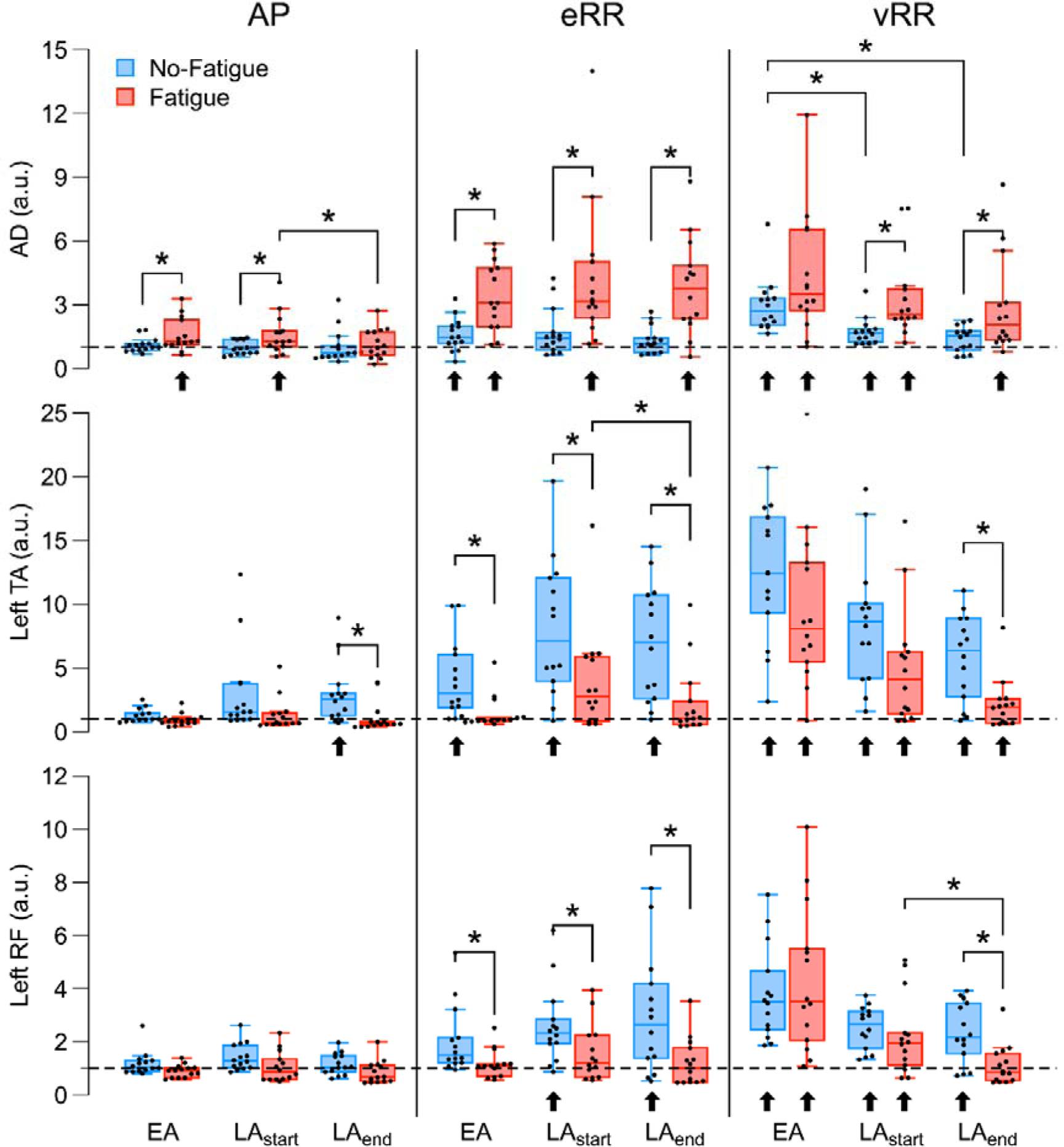
EMG activity of individual muscles for each predetermined time window across the adaptation phase. Muscles are reported on different lines, while EMG time-windows are placed side-by-side in chronological order. Box plots between the 25^th^ and 75^th^ quartiles and the whisker showing the min and max values with median values (thick horizontal lines) are presented. Values are normalized participant-wise to the LB values and reported in arbitrary units (a.u.). * indicates significant between/within group effects (p < 0.05); ⬆ significant difference from LB (p < 0.05). Black horizontal dashed line =1 represents LB values. AD: anterior deltoid, AP: anticipatory phase, BF: biceps femoris, EA: early adaptation, eRR: early Reactive Response, EW: early washout, GM: gastrocnemius medialis, LA_end_: late adaptation – final block, LA_start_: late adaptation – start, LB: late baseline, LW: late washout, PD: posterior deltoid, RF: rectus femoris, TA: tibialis anterior, vRR: voluntary Reactive Response.

**Figure 5.**
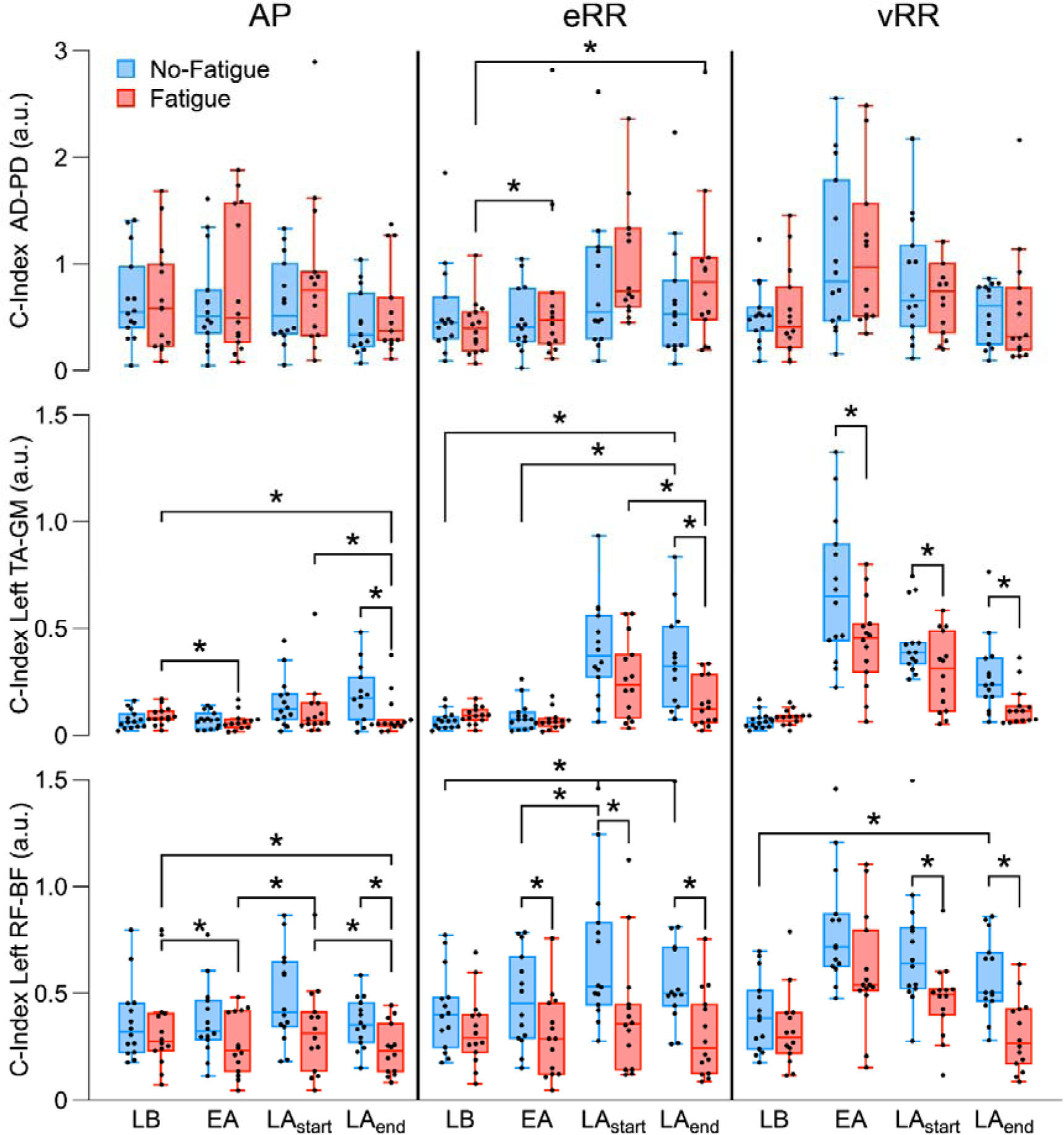
Co-activation index (C-index) at the level of agonist-antagonist muscle pairs across the adaptation phase. Muscle pairs are reported on different lines, while EMG time-windows are placed side-by-side in chronological order. Box plots between the 25^th^ and 75^th^ quartiles and the whisker showing the min and max values with median values (thick horizontal lines) are presented. Values are normalized participant-wise and reported in arbitrary units (a.u.). * indicates significant between/within group effects (p < 0.05). AD: anterior deltoid, AP: anticipatory phase, BF: biceps femoris, EA: early adaptation, eRR: early Reactive Response, EW: early washout, GM: gastrocnemius medialis, LA_end_: late adaptation – final block, LA_start_: late adaptation – start, LB: late baseline, LW: late washout, PD: posterior deltoid, RF: rectus femoris, TA: tibialis anterior, vRR: voluntary Reactive Response.

### Fatiguing exercise produced significant reductions in MVC of the fatigued muscles

In our study, baseline measurements of maximum voluntary contraction (MVC) isometric ankle dorsiflexion force were comparable between the *Fatigue* Group (264.3 ± 78.2 N) and the *No-Fatigue* Group (261.8 ± 86.1 N), indicating no difference in initial strength levels between the groups (p > 0.05). The average duration of the isometric fatiguing exercise was 4 min 15 s (± 2 min,59 s), with an average of 6.6 (± 3.2) cycles. Notably, the duration of exercise decreased across bouts, with the first bout lasting 7 min, 56 s (± 4 min, 14 s) and subsequent bouts showing progressively shorter durations (second bout: 4 min, 4 s ± 1 min, 47 sec, third bout: 3 min, 30 s ± 1 min, 41 s, fourth bout: 3 min, 19 sec ± 1 min, 37 s, fifth bout: 2 min, 27 s ± 1 min, 58 s). Following the fatiguing exercise, a reduction in MVC isometric force was observed exclusively in the Fatigue (−32 ± 4%), compared to the No-Fatigue (1 ± 3%) group, indicating a pronounced effect of the isometric fatiguing exercise on muscle force in the fatigue condition (p < 0.001). Power spectral median frequency decreased significantly in both groups, but the decrease of 26.3% was much bigger in the Fatigue group (*Fatigue* 1^st^ cycle: 112 ± 27.4 Hz – last cycle: 82.5 ± 20.7 Hz, t(_13_) = 5.43, p < 0.001, *d*= 1.45 [C.I: 0.677–2.2]) compared to the 6.7% decrease in the No-Fatigue group (*No-Fatigue* 1^st^ cycle: 119 ± 18 Hz – last cycle: 111 ± 18.2 Hz, t(_13_) = 4.64, p < 0.001, *d*= 1.24 [C.I: 0.522–1.93]).

### Reaching performance exhibited phase dependent adaptive changes and the Fatigue group adapted slower than the No-Fatigue group

We measured reaching movement performance by measuring HME, the peak signed perpendicular deviation of the hand trajectory from a straight line connecting the two targets, and the peak hand velocity in the anteroposterior direction, HPV. Note that since the intended movement was along the medial-lateral direction, we expected HPV to be very small for baseline blocks and return to almost zero values late in the adaptation.

HME and HPV exhibited a decline across both the Fatigue and No-Fatigue groups during the adaptation task, as shown in **Figure 3A**. Additional analyses were performed considering HME and HPV group-averaged data during the first adaptation block (see Methods). The fitting lines described quite well the HME dataset, as indicated by goodness of fit parameters (adj R^2^=0.69 and 0.72; RMSE= 0.34 and 0.3 for Fatigue and No-Fatigue, respectively). Fitting coefficients and respective confidence intervals (C.I. – in square brackets) of the fittings were similar between the groups for terms *a* and *c* (Fatigue: 0.21 [0.13 – 0.29], No-Fatigue: 0.21 [0.14 – 0.29]; Fatigue: 4.03 [3.22 – 4.83], No-Fatigue: 3.48 [2.76 – 4.20], respectively). Coefficient *b* was higher in the Fatigue group (Fatigue: 1.04 [C.I. 0.93 – 1.16], No-Fatigue: 0.65 [C.I. 0.55 – 0.75]), confirming the higher steady-state final value (**Figure 3**, **B** - left side).

Fitting lines for HPV values described quite well the dataset, as indicated by goodness of fit parameters (adj R^2^=0.94 and 0.92; RMSE= 0.01 and 0.01 for Fatigue and No-Fatigue, respectively). Despite a small difference in coefficient *a*, the confidence intervals for the slope of the fittings did not differ between the groups (Fatigue: 0.055 [C.I. 0.042 – 0.067], No-Fatigue: 0.078 [C.I. 0.062 – 0.094]. The other two fitting coefficients, *b* (F: 0.171 [C.I. 0.161 – 0.18], No-Fatigue: 0.151 [C.I. 0.147 – 0.156]) and *c* (Fatigue: 0.307 [C.I. 0.297 – 0.318], No-Fatigue: 0.252 [C.I. 0.242 – 0.262]) were higher for the Fatigue group, denoting both a higher initial value and a higher steady-state final value for HPV in the Fatigue group (**Figure 3**, **B** - right side).

Considering that most of our data did not meet normality assumption, a non-parametric statistical model was employed (i.e. Friedman’s test for repeated measures) to test for factor Phase (*LB*: Late Baseline, *EA*: Early Adaptation, *LA_start_*: Start of Late Adaptation, and *LA_end_*: End of Late Adaptation). We observed a main effect of Phase on HME (χ² (3) = 73.6, p < 0.001), which differed across all pairwise comparisons (p < 0.001). **Figure 3C** illustrates a sharp increase in HME transitioning from *LB* to *EA*, followed by a gradual decrease throughout the adaptation phase.

Conversely, during the washout phase, an initial negative trend in HME was observed, which subsequently approached values near zero towards the end of adaptation. This pattern was statistically confirmed by examining Phase effects, with early washout (EW) and late washout (LW) as specific intervals. A repeated measures ANOVA revealed a main effect for Phase (F_(1.35, 35.22)_ = 194.26, p < 0.001), underscoring the dynamic changes in HME over the course of the study.

To test the effect of Fatigue, we performed independent samples Welch’s t-test to compare between Fatigue and No-Fatigue groups in each Phase. The test showed a difference in HME between groups both at *LA_start_* (Fatigue: 2.06 ± 0.93 cm, No-Fatigue: 1.08 ± 0.98 cm; t_(25.9)_= −2.739, p = 0.011, Cohen’s *d* = 1.035) and at *LA_end_* phases (Fatigue: 0.85 ± 0.77, cm No-Fatigue: 0.17 ± 0.74 cm; t_(25.9)_= −2.385, p = 0.025, *d* = 0.902), where HME was higher in the Fatigue group.

HPV exhibited a pattern throughout the experiment that paralleled HME. The influence of the experimental phase on HPV was significant (χ² (3) = 65.2, p < 0.001), while the reduction in HPV values from the start (*LA_start_*) to the end (*LA_end_*) of the late adaptation phase was significant only for the Fatigue group. Welch’s t-test conducted during the adaptation phase revealed a difference in HPV between the two groups at the start (*LA_start_*) of the late adaptation phase, with the Fatigue group showing higher HPV (0.21 ± 0.08 m/s) compared to the No-Fatigue group (0.16 ± 0.05 m/s; t(_26_)= −2.296, p = 0.003, *d* = 0.87, as illustrated in **Figure 3**, **C**).

Similarly, during the washout phase, HPV continued to follow the trend observed during the adaptation phase. A main effect for Phase was noted (χ² (2) = 42.3, p < 0.001), and a Welch’s t-test identified a difference in HPV between the groups at the early washout (*EW*) phase, with the Fatigue displaying higher HPV values (0.33 ± 0.13 m/s) compared to the No-Fatigue group (0.24 ± 0.09 m/s; t_(22.6)_= −2.1206, p = 0.045, *d* = 0.8, as shown in **Figure 3**, **C**).

### Neuromuscular fatigue induced changes in muscle activity occurred mainly in upper extremity muscles and left side of the lower body

Effects of NMF were more evident on the left-side (contralateral) of the body and effector (i.e. shoulder) muscles. Notable changes in EMG patterns were seen across the adaptation experimental phases (*EA*, *LA_start_* and *LA_end_*; **Figure 4**) in both groups. Differences in muscle activation compared to *LB* phase (p< 0.05) were also present and are illustrated in **Figure 4** and **Supplemental material (S1–S3)** using the arrow symbol below the box plots.

In detail, across each time-window (AP, eRR, vRR) we observed higher *anterior deltoid* (AD) muscle activity in the Fatigue compared to the No-Fatigue group (**Figure 4**), first row; **Table 2**). Compared to baseline (*LB)* values, AD activity increased across all time windows, especially in the Fatigue group (bottom arrow symbols in **Figure 4**, first row). EMG activation of left *tibialis anterior* and *rectus femoris* muscles (**Figure 4**, 2^nd^ and 3^rd^ row; and **Table 2**) generally increased across experimental phases. In contrast to the AD muscle, where the Fatigue group showed higher activation, EMG activation was generally higher in No-Fatigue compared to the Fatigue group for left TA and left RF. Overall, we observed a reduction in EMG activation for each muscle across the experimental phases during vRR. The statistical results are summarized in Table 2. Supplementary results during the *adaptation* phase for the other muscles are reported in Supplementary materials (**S1** and **S2**).

Results for the *washout* phase partially reflect the typical aftereffects seen in the performance variables, with EMG values slowly returning to values close to late baseline (*LB)*. Differences between groups are appreciable in AP, eRR and vRR windows in both upper- and lower-limb muscles.

Overall, the aftereffects were similar to the adaptation phase results. We observed increased AD activation for the Fatigue group, especially during eRR window, while activation of left TA and RF muscles was higher in the No-Fatigue group during the same window. The washout results are summarized in Table 2. Detailed results for the *washout* phase are reported in Supplementary materials (**S1, S2** and **S3**).

Similarly to EMG amplitudes of individual muscles, NMF affected mainly the co-activation index (C-index) at the level of the shoulder and the left leg. Specifically, co-contraction between the anterior deltoids (AD) and posterior deltoids (PD) increased during adaptation, especially in the Fatigue group. During the voluntary Reactive Response (vRR) window C-index increased similarly in both groups following the introduction of the perturbation (*EA*) and later decreased approaching values close to late baseline (*LB*) at the end (*LA_end_*) of the adaptation phase (**Figure 5**, first row).

C-index for left TA-GM and RF-BF muscle pairs was higher in the No-Fatigue group across the adaptation phase during each window. C-index demonstrated an increasing trend across adaptation during both anticipatory phase (AP) and early Reactive Response (eRR) windows, especially for the No-Fatigue group. During vRR, C-index showed a decreasing trend, mainly for the left TA-GM muscle pair (**Figure 5**, 2^nd^ and 3^rd^ row; **Table 3**).

The C-index of right-side muscle pairs increased across the adaptation phase, especially in the No-Fatigue group. Differences between groups, however, were less evident (see Supplementary Materials, **S4**).

Trends for C-index during washout phase were similar to individual EMGs results and demonstrate an after-effect following the removal of the perturbation during early washout (*EW*). Aftereffects in AD-PD muscle pair C-index were significant only for the Fatigue group. The C-index of left TA-GM and RF-BF muscle pairs showed increased values in the No-Fatigue compared to the Fatigue group. C-index similarly decreased in both groups from *EW* to *LW* (see Supplementary Materials, **S5**).

Unlike EMG amplitudes that primarily changed on the left side of the body, C-index of right-side muscle pairs highlighted group differences only for RF-BF at *LW* during the anticipatory phase, and close to significance for *EW*. Overall, the decrease in C-index across the washout phase was significant mainly in the Fatigue group (see Supplementary Materials, **S4**).

### Center-of-pressure excursion is stabilized during neuromuscular fatigue

The displacement of the center of pressure was similar between the two groups (**Figure 6A**). Statistical results for measures of postural stability across experimental phases are shown in **Figure 6B**. Repeated measures ANOVA revealed a main effect of *Phase* on CoP_forward_ (F_(2.03, 52.81)_ = 33.10, p < 0.001, η^2^= 0.393). Holm-Bonferroni post hoc pairwise tests indicated values were different across each condition (p < 0.05) except for *LA_start_*–*LA_end_* comparison. A main effect of *Phase*Group* interaction was also found (F_(2.03, 52.81)_ = 3.48, p = < 0.037, η^2^= 0.041). Holm-Bonferroni post hoc pairwise tests indicated values at EA were higher for No-Fatigue (p = 0.036; **Figure 6**, **B** – top panel).

**Figure 6.**
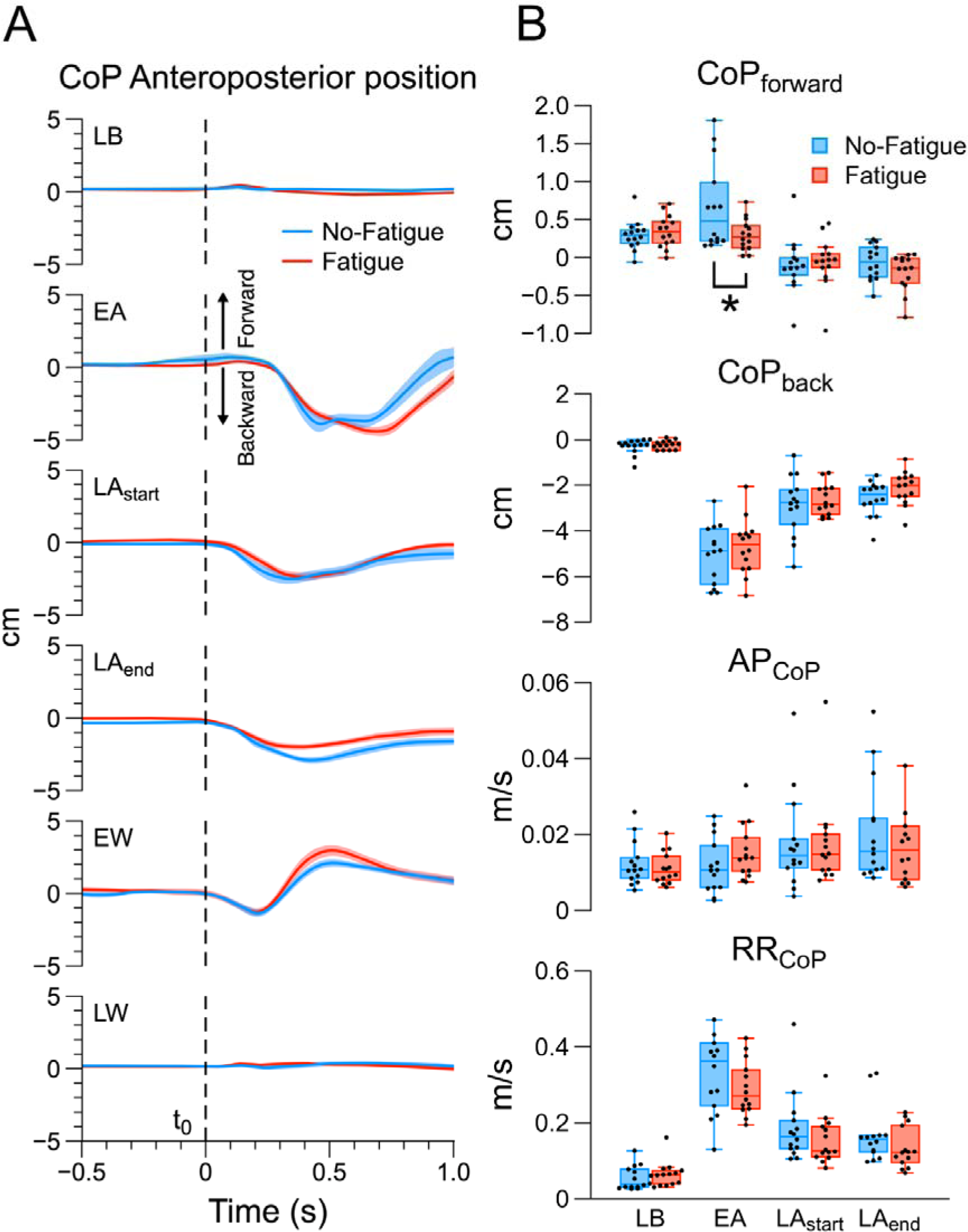
**A:** Group-averaged CoP_y_ trajectory aligned at movement onset (t0) across experimental phases. Thick lines represent means and shaded areas depict ± 1 Standard Error (SE). Vertical dashed lines represent hand movement onset (t0). Forward (positive) and backward (negative) displacements are indicated by the thick black arrows. **B:** Measures of postural stability across the adaptation phase. * indicates significant between/within group effects (p < 0.05). Individual data points (dots and diamonds) with median values (black horizontal lines) are presented. First panel (CoP_forward_): maximal forward displacement of CoP during the reaching movement; Second panel (CoP_back_): maximal backward displacement of CoP during the reaching movement; Third panel (anticipatory phase of CoP – AP_CoP_): mean CoP_y_ velocity during anticipatory phase; Fourth panel (reactive response of CoP – RR_CoP_): peak CoP_y_ velocity during the movement.EA: early adaptation, EW: early washout, LA_end_: late adaptation – final block, LA_start_: late adaptation – start, LB: late baseline, LW: late washout.

A main effect of *Phase* was found for CoP_back_ (F_(2.23, 55.73)_ = 171.95, p = < 0.001, η^2^= 0.766 BF10 =), where Holm-Bonferroni post hoc pairwise comparisons were significant for each condition (p < 0.001; p= 0.012 for *LA_start_*–*LA_end_*). *Phase* effect was found also for AP_CoP_ (F_(1.93, 48.36)_ = 4.7, p= 0.048, η^2^= 0.093), where Holm-Bonferroni post hoc pairwise comparisons revealed differences only between LB–*LA_start_* and LB–*LA_end_* comparisons (LB: 0.0115 ± 0.005: *LA_start_* 0.0177 ± 0.012; *LA_end_*: 0.205 ± 0.017; p= 0.045). Finally, a *Phase* effect was confirmed also for RR_CoP_ (F_(1.9, 47.61)_ = 82.8, p < 0.001, η^2^= 0.589). Holm-Bonferroni post hoc pairwise comparisons were significant for each condition (p < 0.05). Despite the visually appreciable modifications in CoP_y_ profiles across phases (**Figure 6**, **A**), there were no effects of *Group* and *Phase*Group* interaction in both *adaptation* and *washout* phases on measures of postural stability – except for CoP_forward_ at EA (**Figure 6**, **B**). Bayesian rm ANOVA results confirmed the evidence for the equality between groups for all CoP-related variables, where the evidence in favor of the *Phase* model (BF_01_; Bayes’ factor for *H0* over *H1*) – except for CoP_forward_ – were always higher (CoP_back_: *Phase* model compared to *Group* model, BF_01_= 2.804*10^35^ – very strong evidence; AP_CoP_: *Phase* model compared to *Group* model, BF_01_= 35.89 – strong evidence; RR_CoP_: *Phase* model compared to *Group* model, BF_01_= 3.001*10^22^ – very strong evidence). Results for CoP variables during the *washout* phase are reported in Supplementary materials (**S6**).

## Discussion

In this study, we examined how localized neuromuscular fatigue (NMF) in the tibialis anterior, a key postural muscle, influences the adaptive mechanisms necessary for performing human reaching movements while standing upright. We employed a force field perturbation paradigm as our experimental approach to investigate the processes of motor adaptation. Our findings revealed that postural control metrics did not change following exercises that induced fatigue, supporting our initial hypothesis that individuals would prioritize maintaining their posture to aid in performing reaching tasks. This prioritization was achieved through bilateral adjustments in muscle activity, both in muscles subjected to fatiguing exercises and those that were not.

The study also demonstrated that NMF adversely affected the ability to execute reaching movements during both the initial and final phases of adaptation to the force field, identified as LA_start_ and LA_end_ phases, respectively. This result highlighted that fatigue led to greater errors in movement during these phases, though it did not alter the rate of adaptation. Furthermore, we observed differences in the activation patterns of individual muscles and the co-activation patterns of muscle pairs among participants. These differences indicate that the group experiencing fatigue employed unique strategies, likely indicative of a central adaptation mechanism, to mitigate the impact of NMF on the tibialis anterior muscles.

### Postural control following localized neuromuscular fatigue exhibits task prioritization

We monitored both anticipatory and reactive postural adjustments in participants across two cohorts – those experiencing localized neuromuscular fatigue (Fatigue) and those without fatigue (No-Fatigue) – when exposed to force field disturbances. Anticipatory responses, which are preparatory adjustments made before the commencement of reaching tasks, and reactive responses, immediate adjustments in response to postural perturbations, were observed in participants of both groups. This finding is in agreement with earlier studies (41, 53), suggesting a similar adaptation in postural control strategies under similar experimental conditions.

Significantly, for both groups, dynamic changes in postural stability metrics occurred over time, particularly in terms of the Center of Pressure’s (CoP) displacement or velocity in the anteroposterior direction, as illustrated in **Figure 6**. Despite these temporal changes, we did not observe significant differences in postural stability between the Fatigue and No-Fatigue groups. Over time, there was a notable reduction in the reactive response of the CoP (RR_CoP_), indicating improvements in online postural corrections, alongside an increase in anticipatory postural control of the CoP (AP_CoP_). This suggests that participants in both groups efficiently developed motor strategies to preemptively counteract and adapt to the disturbances caused by the force field perturbation, enhancing postural stability.

These adaptive behavioral responses are supported by findings from studies that examined the impact of intentional, specific movements of the upper limbs on postural stability (4–6). For example, Strang and colleagues documented modifications in the timing of muscle activation in response to neuromuscular fatigue, affecting both fatigued and non-fatigued muscles (7). Such adjustments in muscle activation demonstrate the central nervous system’s (CNS) involvement, particularly supraspinal structures, in modifying neural signals to various muscles. The results from our study suggest that the purpose of this adjustment is to preserve postural stability during the performance of tasks that require precision with the upper limbs, underscoring the CNS’s critical function in orchestrating and harmonizing motor responses across the body by prioritizing control of posture.

The findings of Strang and colleagues are congruent with our observations concerning muscle activation patterns (7). Despite identifying differences in activation at both the level of individual muscles (**Figure 4**) and in the coordinated activation of muscle pairs (**Figure 5**), we observed no alterations in postural stability following fatiguing exercises. This finding is echoed in the work of Singh and Latash, who also noted no significant shifts in the CoP during voluntary sway tasks performed while standing, even though fatigue had modified patterns of muscle activation and muscle synergies (17). Furthermore, a more recent study revealed that although exercises targeting the soleus muscle altered EMG activity, they did not affect balance performance in dynamic postural tasks (26). These outcomes suggest that in the context of complex motor tasks, such as maintaining an upright posture which necessitates the involvement of multiple joints and muscles (42, 72), the CNS implements strategies to ensure the stabilization of motor performance during fatigue (7).

Prioritization of postural balance during reaching movements has been substantiated through perturbation tasks. Trivedi et al. investigated the effects of unexpected perturbations of the standing surface and found that compensatory postural responses were modulated to facilitate reaching movements (73). Their findings indicate that long-latency components of compensatory postural responses in the lower extremities adapt to ensure stable reaching performance. Similarly, Pienciak-Siewert et al. demonstrated a “posture first” strategy by applying a velocity-dependent force field, creating distinct mechanical perturbations in forward and backward directions (54). Despite similar reaching adaptation in both directions, they observed direction-specific coactivation of postural muscles. These results further support the notion that the CNS prioritizes postural control and movement adaptation to achieve complex task objectives that require upright posture stabilization.

Both force-field adaptation and the long-latency components of reflexes are mediated by the cerebellum, suggesting its pivotal role in establishing a task prioritization hierarchy to stabilize motor performance during upper limb tasks performed in an upright stance. This hypothesis is supported by a cerebellar lesion study conducted by Ilg et al. In their dual-task experiment, which involved both working memory and walking, patients with focal cerebellar lesions exhibited impaired performance on the cognitive task but not on the walking task, despite displaying signs of ataxia (74). These findings indicate that, even with motor impairments due to focal cerebellar injury, the unaffected regions of the cerebellum prioritize motor performance and balance over cognitive task execution.

#### Fatigue-induced effects on reaching performance in a force field

The performance in reaching tasks, quantified by hand movement error (HME) – the maximum deviation of the hand’s path from a straight line between targets – demonstrated a trend of exponential decrease, as illustrated in **Figure 3A**. This observation is consistent with findings from similar research on motor adaptation in reaching tasks conducted in a standing position (41, 53, 54).

In this study, the impact of neuromuscular fatigue on hand movement error (HME) and hand peak velocity (HPV) were significant across some experimental phases. Although the adaptation phase showed a reduction in HME and HPV comparable to that seen in non-fatigued conditions (**Figure 3**, **A**), and similar patterns of aftereffects were observed after perturbation removal (washout phase), the Fatigue group experienced an altered adaptation process for both the variables. This was evident from elevated HME and HPV values at the conclusion of both the first block and the entire adaptation phase (**Figure 3**, **C**).

It has been proposed that the fast and slow learning phases correspond to explicit and implicit learning processes, respectively (76, 77). From a computational perspective, explicit learning is geared towards achieving specific goals, whereas implicit learning focuses on the precise execution of movements (78). This distinction is mirrored in the underlying neural mechanisms: strategies for target acquisition, which are employed to counter experimentally induced disturbances, are governed by prefrontal cortex activity. In contrast, implicit learning depends on the cerebellum (79–84), with proprioception playing a key role in driving implicit learning (85, 86).

Our findings diverge to some extent from the results of Takahashi et al. (40), who reported no impact of NMF on motor adaptation in reaching tasks. This discrepancy may stem from differences in experimental design. Specifically, Takahashi et al.’s participants initially adapted to the reaching task without fatigue, potentially achieving performance stabilization before encountering NMF. Additionally, their adaptation phase was limited to 20 trials, possibly too short for highlighting long-term differences. Another possible explanation might arise from the different fatiguing protocols between the study by Takahashi and the present one. In their study they fatigued participants’ effector muscles (i.e. muscles in the upper limb) during resisted reaching movements, which possibly allowed the CNS to process and integrate proprioceptive signals coming from the fatigued muscles (i.e. group III and IV afferences (87, 88)) prior to the following execution of the retention tests.

### Fatigue-induced changes in EMG activation and co-activation patterns

EMG analysis revealed divergent activation patterns in the musculature of the upper and lower limbs between subjects in the Fatigue and No-Fatigue groups throughout the experimental stages. Specifically, individuals in the Fatigue group exhibited higher levels of muscle activity in the upper extremity, alongside lower levels of activity in the leg muscles, as depicted in **Figure 4**. Notably, differences in EMG activation were primarily observed in postural muscles on the contralateral side with respect to the reaching arm. This observation aligns with the mechanism where, during a directional reaching task, postural muscles on the body’s contralateral side with respect to target direction are activated earlier than ipsilateral muscle and even before the onset of the reaching action (89). This preemptive activation is likely related to the biomechanical constraints of the reaching movement (i.e. moving the hand from midline to the right requires a preparatory shifting of CoP in the medio-lateral direction) and serves to stabilize posture in anticipation of a volitional movement towards the target. Our findings provide support for studies that have shown feedforward modulation of lower limb muscle activity in relation to the direction of movement (89–91).

The observed decrease in lower-limb muscle activation, particularly during the anticipatory phase, in the Fatigue group implies that fatigue affects the pre-programming of movements involving postural muscles. Leonard and colleagues suggested that anticipatory activity in postural muscles contralateral to the reaching arm is evidence of CNS mechanisms at play (89). The diminished or absent anticipatory activity in both the fatigued muscle (e.g., tibialis anterior) and the non-fatigued muscle (e.g., rectus femoris) in the Fatigue group indicates that the fatiguing exercise exerted a widespread impact, affecting motor planning at the central level. This finding aligns with prior studies on the effects of localized fatigue on motor control (17, 23) and, to our understanding, marks a clear demonstration of such neural drive reorganization during motor adaptation.

Conversely, the Fatigue group also exhibited increased activation of the anterior deltoid (AD) muscle, alongside more frequent adjustments in hand movement trajectory – as evidenced by higher hand path variability (HPV) values (**Figure 3**) – suggesting a greater reliance on visual input for correcting movement trajectories. This may be an attempt to offset the unreliable proprioceptive feedback from fatigued muscles (14, 88, 92). Additionally, the elevated AD muscle activity during the anticipatory phase suggests that participants pre-activated this muscle to enhance its response to force field disturbances, thereby increasing the muscle’s “apparent stiffness”, which has been broadly defined as the change in force per unit change in joint angle (93, 94).

In conclusion, our findings indicate that localized neuromuscular fatigue adversely affects the adaptation of reaching movements in response to force field perturbations. This impairment does not stem from a reduction in postural stability, which remains intact following NMF, but is likely due to compromised sensorimotor integration, particularly in processing proprioceptive and sensory feedback during fatigue. Notably, the cerebellum’s critical role in motor adaptation may underlie these observed discrepancies. Our study also reveals variations in the activation of both fatigued and non-fatigued muscles, as well as in the strategies for correcting movements, indicating that the physiological disruptions caused by NMF during force field perturbations could prompt the selection of alternative motor adaptation strategies. Future research should investigate whether these adaptations are temporary, resolving with recovery, or if they have lasting impacts on the retention of these newly adopted strategies.

## SUPPLEMENTAL MATERIAL

Available at OSF: https://osf.io/hgpu9/

**S1** – Upper limb & right-side muscles EMGs (Text, Figures & Tables)

**S2** – Dorsal EMGs (Text, Figures & Tables)

**S3** – Left EMGs washout (Figure)

**S4** – Right C-index (Figures & Tables)

**S5** – Left C-index washout (Figure)

**S6** – CoP variables washout (Figure)

**S7** – Results for the correlation analysis between performance variables and stability measures

## References

1. Bigland-Ritchie B, Furbush F, Woods JJ. Fatigue of intermittent submaximal voluntary contractions: central and peripheral factors. J Appl Physiol 61: 421–429, 1986. doi: 10.1152/jappl.1986.61.2.421.

2. Gandevia SC. Spinal and supraspinal factors in human muscle fatigue. Physiol Rev 81: 1725–1789, 2001. doi: 10.1152/physrev.2001.81.4.1725.

3. Enoka RM, Duchateau J. Muscle fatigue: what, why and how it influences muscle function: Muscle fatigue. J Physiol 586: 11–23, 2008. doi: 10.1113/jphysiol.2007.139477.

4. Strang AJ, Berg WP. Fatigue-induced adaptive changes of anticipatory postural adjustments. Exp Brain Res 178: 49–61, 2007. doi: 10.1007/s00221-006-0710-5.

5. Gates DH, Dingwell JB. The effects of neuromuscular fatigue on task performance during repetitive goal-directed movements. Exp Brain Res 187: 573–585, 2008.

6. Kanekar N, Santos MJ, Aruin AS. Anticipatory postural control following fatigue of postural and focal muscles. Clin Neurophysiol 119: 2304–2313, 2008. doi: 10.1016/j.clinph.2008.06.015.

7. Strang AJ, Berg WP, Hieronymus M. Fatigue-induced early onset of anticipatory postural adjustments in non-fatigued muscles: support for a centrally mediated adaptation. Exp Brain Res 197: 245–254, 2009. doi: 10.1007/s00221-009-1908-0.

8. Forestier N, Nougier V. The effects of muscular fatigue on the coordination of a multijoint movement in human. Neurosci Lett 252: 187–190, 1998. doi: 10.1016/S0304-3940(98)00584-9.

9. Coventry E, O’Connor KM, Hart BA, Earl JE, Ebersole KT. The effect of lower extremity fatigue on shock attenuation during single-leg landing. Clin Biomech 21: 1090–1097, 2006. doi: 10.1016/j.clinbiomech.2006.07.004.

10. Singh T, SKM V, Zatsiorsky VM, Latash ML. Fatigue and motor redundancy: adaptive increase in finger force variance in multi-finger tasks. J Neurophysiol 103: 2990–3000, 2010.

11. Singh T, Zatsiorsky VM, Latash ML. Contrasting effects of fatigue on multi-finger coordination in young and older adults. J Appl Physiol 115: 456–467, 2013.

12. Nardone A, Tarantola J, Giordano A, Schieppati M. Fatigue effects on body balance. Electroencephalogr Clin Neurophysiol 105: 309–320, 1997. doi: 10.1016/s0924-980x(97)00040-4.

13. Vuillerme N, Boisgontier M. Muscle fatigue degrades force sense at the ankle joint. Gait Posture 28: 521–524, 2008. doi: 10.1016/j.gaitpost.2008.03.005.

14. Albanese GA, Falzarano V, Holmes MWR, Morasso P, Zenzeri J. A Dynamic Submaximal Fatigue Protocol Alters Wrist Biomechanical Properties and Proprioception. Front Hum Neurosci 16: 887270, 2022. doi: 10.3389/fnhum.2022.887270.

15. Vafadar AK, Côté J, Archambault PS. The effect of muscle fatigue on position sense in an upper limb multi-joint task. Motor Control 16: 265–283, 2012.

16. Grose G, Manzone DM, Eschelmuller G, Peters RM, Carpenter MG, Inglis JT, Chua R. The effects of eccentric exercise-induced fatigue on position sense during goal-directed movement. J Appl Physiol 132: 1005–1019, 2022. doi: 10.1152/japplphysiol.00177.2021.

17. Singh T, Latash ML. Effects of muscle fatigue on multi-muscle synergies. Exp Brain Res 214: 335–350, 2011. doi: 10.1007/s00221-011-2831-8.

18. Segers V, Lenoir M, Aerts P, De Clercq D. Influence of M. tibialis anterior fatigue on the walk-to-run and run-to-walk transition in non-steady state locomotion. Gait Posture 25: 639–647, 2007. doi: 10.1016/j.gaitpost.2006.07.008.

19. Parijat P, Lockhart TE. Effects of quadriceps fatigue on the biomechanics of gait and slip propensity. Gait Posture 28: 568–573, 2008. doi: 10.1016/j.gaitpost.2008.04.001.

20. Qu X, Yeo JC. Effects of load carriage and fatigue on gait characteristics. J Biomech 44: 1259–1263, 2011. doi: 10.1016/j.jbiomech.2011.02.016.

21. Barbieri FA, dos Santos PCR, Vitório R, van Dieën JH, Gobbi LTB. Effect of muscle fatigue and physical activity level in motor control of the gait of young adults. Gait Posture 38: 702–707, 2013. doi: 10.1016/j.gaitpost.2013.03.006.

22. Gribble PA, Hertel J. Effect of lower-extremity muscle fatigue on postural control. Arch Phys Med Rehabil 85: 589–592, 2004. doi: 10.1016/j.apmr.2003.06.031.

23. Lyu H, Fan Y, Hao Z, Wang J. Effect of local and general fatiguing exercises on disturbed and static postural control. J Electromyogr Kinesiol 56: 102487, 2021. doi: 10.1016/j.jelekin.2020.102487.

24. Lyu H, Fan Y, Hua A, Cao X, Gao Y, Wang J. Effects of unilateral and bilateral lower extremity fatiguing exercises on postural control during quiet stance and self-initiated perturbation. Hum Mov Sci 81: 102911, 2022. doi: 10.1016/j.humov.2021.102911.

25. Davidson BS, Madigan ML, Nussbaum MA, Wojcik LA. Effects of localized muscle fatigue on recovery from a postural perturbation without stepping. Gait Posture 29: 552–557, 2009. doi: 10.1016/j.gaitpost.2008.12.011.

26. Marcolin G, Cogliati M, Cudicio A, Negro F, Tonin R, Orizio C, Paoli A. Neuromuscular Fatigue Affects Calf Muscle Activation Strategies, but Not Dynamic Postural Balance Control in Healthy Young Adults. Front Physiol 13: 799565, 2022. doi: 10.3389/fphys.2022.799565.

27. Paillard T, Chaubet V, Maitre J, Dumitrescu M, Borel L. Disturbance of contralateral unipedal postural control after stimulated and voluntary contractions of the ipsilateral limb. Neurosci Res 68: 301–306, 2010. doi: 10.1016/j.neures.2010.08.004.

28. Singh T, Zatsiorsky VM, Latash ML. Effects of fatigue on synergies in a hierarchical system. Hum Mov Sci 31: 1379–1398, 2012.

29. Halperin I, Chapman DW, Behm DG. Non-local muscle fatigue: effects and possible mechanisms. Eur J Appl Physiol 115: 2031–2048, 2015. doi: 10.1007/s00421-015-3249-y.

30. Casamento-Moran A, Mooney RA, Chib VS, Celnik PA. Cerebellar excitability regulates physical fatigue perception. J Neurosci 43: 3094–3106, 2023.

31. Debaere F, Wenderoth N, Sunaert S, Van Hecke P, Swinnen SP. Cerebellar and premotor function in bimanual coordination: parametric neural responses to spatiotemporal complexity and cycling frequency. Neuroimage 21: 1416–1427, 2004.

32. Soteropoulos DS, Baker SN. Bilateral representation in the deep cerebellar nuclei. J Physiol 586: 1117–1136, 2008.

33. Debaere F, Swinnen SP, Béatse E, Sunaert S, Van Hecke P, Duysens J. Brain areas involved in interlimb coordination: A distributed network. NeuroImage 14: 947–958, 2001. doi: 10.1006/nimg.2001.0892.

34. Maschke M, Gomez CM, Ebner TJ, Konczak J. Hereditary cerebellar ataxia progressively impairs force adaptation during goal-directed arm movements. J Neurophysiol 91: 230–238, 2004.

35. Smith MA, Shadmehr R. Intact ability to learn internal models of arm dynamics in Huntington’s disease but not cerebellar degeneration. J Neurophysiol 93: 2809–2821, 2005.

36. Criscimagna-Hemminger SE, Bastian AJ, Shadmehr R. Size of error affects cerebellar contributions to motor learning. J Neurophysiol 103: 2275–2284, 2010.

37. Donchin O, Rabe K, Diedrichsen J, Lally N, Schoch B, Gizewski ER, Timmann D. Cerebellar regions involved in adaptation to force field and visuomotor perturbation. J Neurophysiol 107: 134–147, 2012.

38. Statton MA, Vazquez A, Morton SM, Vasudevan EVL, Bastian AJ. Making sense of cerebellar contributions to perceptual and motor adaptation. The Cerebellum 17: 111–121, 2018. doi: 10.1007/s12311-017-0879-0.

39. Shadmehr R, Mussa-Ivaldi FA. Adaptive representation of dynamics during learning of a motor task. J Neurosci 14: 3208–3224, 1994.

40. Takahashi CD, Nemet D, Rose-Gottron CM, Larson JK, Cooper DM, Reinkensmeyer DJ. Effect of muscle fatigue on internal model formation and retention during reaching with the arm. J Appl Physiol 100: 695–706, 2006. doi: 10.1152/japplphysiol.00140.2005.

41. Ahmed AA, Wolpert DM. Transfer of dynamic learning across postures. J Neurophysiol 102: 2816–2824, 2009. doi: 10.1152/jn.00532.2009.

42. Paillard T. Effects of general and local fatigue on postural control: A review. Neurosci Biobehav Rev 36: 162–176, 2012. doi: 10.1016/j.neubiorev.2011.05.009.

43. Berger LL, Regueme SC, Forestier N. Unilateral lower limb muscle fatigue induces bilateral effects on undisturbed stance and muscle EMG activities. J Electromyogr Kinesiol 20: 947–952, 2010. doi: 10.1016/j.jelekin.2009.09.006.

44. Taylor JL, Butler JE, Gandevia SC. Changes in muscle afferents, motoneurons and motor drive during muscle fatigue. Eur J Appl Physiol 83: 106–115, 2000. doi: 10.1007/s004210000269.

45. Liu JZ, Shan ZY, Zhang LD, Sahgal V, Brown RW, Yue GH. Human Brain Activation During Sustained and Intermittent Submaximal Fatigue Muscle Contractions: An fMRI Study. J Neurophysiol 90: 300–312, 2003. doi: 10.1152/jn.00821.2002.

46. van Duinen H, Renken R, Maurits N, Zijdewind I. Effects of motor fatigue on human brain activity, an fMRI study. NeuroImage 35: 1438–1449, 2007. doi: 10.1016/j.neuroimage.2007.02.008.

47. Tanaka M, Watanabe Y. Supraspinal regulation of physical fatigue. Neurosci Biobehav Rev 36: 727–734, 2012. doi: 10.1016/j.neubiorev.2011.10.004.

48. Takakusaki K, Takahashi M, Obara K, Chiba R. Neural substrates involved in the control of posture. Adv Robot 31: 2–23, 2017. doi: 10.1080/01691864.2016.1252690.

49. Hermens HJ, Freriks B, Disselhorst-Klug C, Rau G. Development of recommendations for SEMG sensors and sensor placement procedures. J Electromyogr Kinesiol 10: 361–374, 2000.

50. Ricotta JM, De SD, Nardon M, Benamati A, Latash ML. Effects of fatigue on intramuscle force-stabilizing synergies. J Appl Physiol Bethesda Md 1985 135: 1023–1035, 2023. doi: 10.1152/japplphysiol.00419.2023.

51. Carroll TJ, Taylor JL, Gandevia SC. Recovery of central and peripheral neuromuscular fatigue after exercise. J Appl Physiol 122: 1068–1076, 2017.

52. Enoka RM, Stuart DG. Neurobiology of muscle fatigue. J Appl Physiol 72: 1631–1648, 1992.

53. Pienciak-Siewert A, Horan DP, Ahmed AA. Trial-to-trial adaptation in control of arm reaching and standing posture. J Neurophysiol 116: 2936–2949, 2016. doi: 10.1152/jn.00537.2016.

54. Pienciak-Siewert A, Horan DP, Ahmed AA. Role of muscle coactivation in adaptation of standing posture during arm reaching. J Neurophysiol 123: 529– 547, 2020. doi: 10.1152/jn.00939.2017.

55. Sinha O, Madarshahian S, Gómez-Granados A, Paine ML, Kurtzer I, Singh T. Smooth pursuit eye movements contribute to anticipatory force control during mechanical stopping of moving objects. J Neurophysiol 129: 1293–1309, 2023. doi: 10.1152/jn.00075.2023.

56. Malone LA, Vasudevan EVL, Bastian AJ. Motor Adaptation Training for Faster Relearning. J Neurosci 31: 15136–15143, 2011. doi: 10.1523/JNEUROSCI.1367-11.2011.

57. Winter DA. Biomechanics and motor control of human movement. 4th ed. Hoboken, N.J: Wiley, 2009.

58. Horak FB, Nashner LM. Central programming of postural movements: adaptation to altered support-surface configurations. J Neurophysiol 55: 1369– 1381, 1986. doi: 10.1152/jn.1986.55.6.1369.

59. Krishnan V, Kanekar N, Aruin AS. Anticipatory postural adjustments in individuals with multiple sclerosis. Neurosci Lett 506: 256–260, 2012. doi: 10.1016/j.neulet.2011.11.018.

60. Scott SH, Cluff T, Lowrey CR, Takei T. Feedback control during voluntary motor actions. Curr Opin Neurobiol 33: 85–94, 2015. doi: 10.1016/j.conb.2015.03.006.

61. Nardone A, Schieppati M. Postural adjustments associated with voluntary contraction of leg muscles in standing man. Exp Brain Res 69: 469–480, 1988. doi: 10.1007/BF00247301.

62. Slijper H, Latash M. The effects of instability and additional hand support on anticipatory postural adjustments in leg, trunk, and arm muscles during standing. Exp Brain Res 135: 81–93, 2000. doi: 10.1007/s002210000492.

63. Slijper H, Latash ML. The effects of muscle vibration on anticipatory postural adjustments. Brain Res 1015: 57–72, 2004. doi: 10.1016/j.brainres.2004.04.054.

64. Piscitelli D, Falaki A, Solnik S, Latash ML. Anticipatory postural adjustments and anticipatory synergy adjustments: Preparing to a postural perturbation with predictable and unpredictable direction. Exp Brain Res 235: 713–730, 2017. doi: 10.1007/s00221-016-4835-x.

65. Bertucco M, Nardello F, Magris R, Cesari P, Latash ML. Postural Adjustments during Interactions with an Active Partner. Neuroscience 463: 14– 29, 2021. doi: 10.1016/j.neuroscience.2021.03.020.

66. Cesari P, Piscitelli F, Pascucci F, Bertucco M. Postural Threat Influences the Coupling Between Anticipatory and Compensatory Postural Adjustments in Response to an External Perturbation. Neuroscience 490: 25–35, 2022. doi: 10.1016/j.neuroscience.2022.03.005.

67. Mathew J, Lefèvre P, Crevecoeur F. Savings in Human Force Field Learning Supported by Feedback Adaptation. eNeuro 8, 2021. doi: 10.1523/ENEURO.0088-21.2021.

68. Wagenmakers E-J, Marsman M, Jamil T, Ly A, Verhagen J, Love J, Selker R, Gronau QF, Šmíra M, Epskamp S, Matzke D, Rouder JN, Morey RD. Bayesian inference for psychology. Part I: Theoretical advantages and practical ramifications. Psychon Bull Rev 25: 35–57, 2018. doi: 10.3758/s13423-017-1343-3.

69. Wagenmakers E-J, Love J, Marsman M, Jamil T, Ly A, Verhagen J, Selker R, Gronau QF, Dropmann D, Boutin B, Meerhoff F, Knight P, Raj A, van Kesteren E-J, van Doorn J, Šmíra M, Epskamp S, Etz A, Matzke D, de Jong T, van den Bergh D, Sarafoglou A, Steingroever H, Derks K, Rouder JN, Morey RD. Bayesian inference for psychology. Part II: Example applications with JASP. Psychon Bull Rev 25: 58–76, 2018. doi: 10.3758/s13423-017-1323-7.

70. Quintana DS, Williams DR. Bayesian alternatives for common null-hypothesis significance tests in psychiatry: a non-technical guide using JASP. BMC Psychiatry 18: 178, 2018. doi: 10.1186/s12888-018-1761-4.

71. Kass RE, Raftery AE. Bayes Factors. J Am Stat Assoc 90: 773–795, 1995. doi: 10.1080/01621459.1995.10476572.

72. Zemková E, Hamar D. Physiological mechanisms of post-exercise balance impairment. Sports Med Auckl NZ 44: 437–448, 2014. doi: 10.1007/s40279-013-0129-7.

73. Trivedi H, Leonard JA, Ting LH, Stapley PJ. Postural responses to unexpected perturbations of balance during reaching. Exp Brain Res 202: 485– 491, 2010. doi: 10.1007/s00221-009-2135-4.

74. Ilg W, Christensen A, Mueller OM, Goericke SL, Giese MA, Timmann D. Effects of cerebellar lesions on working memory interacting with motor tasks of different complexities. J Neurophysiol 110: 2337–2349, 2013. doi: 10.1152/jn.00062.2013.

75. Carroll TJ, Taylor JL, Gandevia SC. Recovery of central and peripheral neuromuscular fatigue after exercise. J Appl Physiol 122: 1068–1076, 2017. doi: 10.1152/japplphysiol.00775.2016.

76. Smith MA, Ghazizadeh A, Shadmehr R. Interacting Adaptive Processes with Different Timescales Underlie Short-Term Motor Learning. PLOS Biol 4: e179, 2006. doi: 10.1371/journal.pbio.0040179.

77. McDougle SD, Bond KM, Taylor JA. Explicit and Implicit Processes Constitute the Fast and Slow Processes of Sensorimotor Learning. J Neurosci Off J Soc Neurosci 35: 9568–9579, 2015. doi: 10.1523/JNEUROSCI.5061-14.2015.

78. Taylor JA, Krakauer JW, Ivry RB. Explicit and Implicit Contributions to Learning in a Sensorimotor Adaptation Task. J Neurosci 34: 3023–3032, 2014. doi: 10.1523/JNEUROSCI.3619-13.2014.

79. Butcher PA, Ivry RB, Kuo S-H, Rydz D, Krakauer JW, Taylor JA. The cerebellum does more than sensory prediction error-based learning in sensorimotor adaptation tasks. J Neurophysiol 118: 1622–1636, 2017. doi: 10.1152/jn.00451.2017.

80. Haar S, Donchin O. A Revised Computational Neuroanatomy for Motor Control. J Cogn Neurosci 32: 1823–1836, 2020. doi: 10.1162/jocn_a_01602.

81. Izawa J, Criscimagna-Hemminger SE, Shadmehr R. Cerebellar Contributions to Reach Adaptation and Learning Sensory Consequences of Action. J Neurosci 32: 4230–4239, 2012. doi: 10.1523/JNEUROSCI.6353-11.2012.

82. Schlerf JE, Xu J, Klemfuss NM, Griffiths TL, Ivry RB. Individuals with cerebellar degeneration show similar adaptation deficits with large and small visuomotor errors. J Neurophysiol 109: 1164–1173, 2013. doi: 10.1152/jn.00654.2011.

83. Tseng Y-W, Diedrichsen J, Krakauer JW, Shadmehr R, Bastian AJ. Sensory prediction errors drive cerebellum-dependent adaptation of reaching. J Neurophysiol 98: 54–62, 2007. doi: 10.1152/jn.00266.2007.

84. Taylor JA, Ivry RB. Cerebellar and prefrontal cortex contributions to adaptation, strategies, and reinforcement learning. Prog Brain Res 210: 217–253, 2014. doi: 10.1016/B978-0-444-63356-9.00009-1.

85. Weeks HM, Therrien AS, Bastian AJ. The cerebellum contributes to proprioception during motion. J Neurophysiol 118: 693–702, 2017. doi: 10.1152/jn.00417.2016.

86. Tsay JS, Kim H, Haith AM, Ivry RB. Understanding implicit sensorimotor adaptation as a process of proprioceptive re-alignment. eLife 11: e76639, 2022. doi: 10.7554/eLife.76639.

87. Amann M, Blain GM, Proctor LT, Sebranek JJ, Pegelow DF, Dempsey JA. Group III and IV muscle afferents contribute to ventilatory and cardiovascular response to rhythmic exercise in humans. J Appl Physiol 109: 966–976, 2010. doi: 10.1152/japplphysiol.00462.2010.

88. Sidhu SK, Weavil JC, Mangum TS, Jessop JE, Richardson RS, Morgan DE, Amann M. Group III/IV locomotor muscle afferents alter motor cortical and corticospinal excitability and promote central fatigue during cycling exercise. Clin Neurophysiol 128: 44–55, 1AD. doi: 10.1016/j.clinph.2016.10.008.

89. Leonard JA, Gritsenko V, Ouckama R, Stapley PJ. Postural adjustments for online corrections of arm movements in standing humans. J Neurophysiol 105: 2375–2388, 2011. doi: 10.1152/jn.00944.2010.

90. Leonard JA, Brown RH, Stapley PJ. Reaching to Multiple Targets When Standing: The Spatial Organization of Feedforward Postural Adjustments. J Neurophysiol 101: 2120–2133, 2009. doi: 10.1152/jn.91135.2008.

91. Lowrey CR, Nashed JY, Scott SH. Rapid and flexible whole body postural responses are evoked from perturbations to the upper limb during goal-directed reaching. J Neurophysiol 117: 1070–1083, 2017. doi: 10.1152/jn.01004.2015.

92. Mohapatra S, Krishnan V, Aruin AS. Postural control in response to an external perturbation: effect of altered proprioceptive information. Exp Brain Res 217: 197–208, 2012. doi: 10.1007/s00221-011-2986-3.

93. Latash ML, Zatsiorsky VM. Joint stiffness: Myth or reality? Hum Mov Sci 12: 653–692, 1993. doi: 10.1016/0167-9457(93)90010-M.

94. Ambike S, Paclet F, Zatsiorsky VM, Latash ML. Factors affecting grip force: anatomy, mechanics, and referent configurations. Exp Brain Res 232: 1219– 1231, 2014. doi: 10.1007/s00221-014-3838-8.

